# Low affinity DNA-binding promotes cooperative activation of natural transformation in *Vibrio cholerae*

**DOI:** 10.64898/2026.01.21.700895

**Authors:** Allison C. Hullinger, Virginia E. Callahan, Triana N. Dalia, Ankur B. Dalia

## Abstract

DNA-binding transcriptional regulators control gene expression in response to environmental cues. A subset of these proteins, called transmembrane transcriptional regulators (TTRs), directly bind DNA to regulate transcription while remaining anchored in the cytoplasmic membrane. Prior work has shown that in the presence of the polysaccharide chitin, two TTRs, TfoS and ChiS, coordinate to induce the expression of TfoR, a small RNA that is critical for natural transformation in *Vibrio cholerae*. Specifically, it was shown that ChiS recruits the P_*tfoR*_ locus to the membrane, which allows for the subsequent activation of this promoter by TfoS. However, it was also shown that increasing TfoS protein levels bypasses this coordination, allowing TfoS to activate the promoter independently. It therefore remains unclear what molecular mechanisms drive the requirement for ChiS in native conditions. Here, we show that ChiS binds P_*tfoR*_ with a higher affinity than TfoS. We hypothesized that the low affinity of TfoS for P_*tfoR*_ helps reinforce its dependence on ChiS for activation. To test this, we isolated a mutant allele of the TfoS DNA-binding domain that has a higher affinity for P_*tfoR*_. We show that this high-affinity TfoS allele promotes ChiS-independent activation of P_*tfoR*_ and dysregulates chitin-dependent phenotypes in *V. cholerae*. These results demonstrate that the relative DNA-binding affinity of these TTRs facilitates their coordination, and that this coordination is necessary for optimal *V. cholerae* fitness on chitin.

**IMPORTANCE:** DNA-binding transmembrane transcriptional regulators (TTRs) are critical for some bacterial species to properly sense and respond to their environments. Recent work highlights that pairs of TTRs can coordinate their activities to regulate gene expression, allowing them to sensitively control behaviors like virulence and horizontal gene transfer. However, the mechanisms that enable this coordination remain poorly understood. Here, we show that the relative DNA-binding affinity of paired TTRs is a critical feature that can drive their coordination.

## INTRODUCTION

DNA-binding transcriptional regulators are a broad class of proteins that directly regulate gene expression in response to environmental stimuli. Transmembrane transcriptional regulators (TTRs) are a subset of DNA-binding regulators that directly regulate expression while remaining anchored in the cytoplasmic membrane (1). Studies performed using the facultative human pathogen *Vibrio cholerae* reveal that multiple TTRs can cooperate to activate the expression of their downstream targets. Specifically, we recently showed that two chitin-induced TTRs in *V. cholerae*, ChiS and TfoS, coordinate to activate the genes necessary for horizontal gene transfer by natural transformation (2). In addition, prior work demonstrates that the TTRs ToxR and TcpP similarly coordinate to activate the expression of the virulence cascade in *V. cholerae* (3-7). In both cases, the cooperative TTRs bind at the same promoter to activate gene expression: ChiS/TfoS bind P_*tfoR*_ (8-10) while ToxR/TcpP bind P_*toxT*_ (6, 11, 12). These examples suggest that coordination may be a common feature of TTR-dependent regulation. Because this cooperative activation relies on the activity of two independent TTRs, this regulatory architecture may allow for tighter regulation of the target genes (*e*.*g*., as a coincidence detector, requiring induction of both TTRs for activation to occur).

When ChiS/TfoS and ToxR/TcpP are expressed at native levels, activation of their target promoters requires both TTR partners. Overexpression of one TTR, however, is sufficient to drive activation in each system: TfoS overexpression is sufficient to activate P_*tfoR*_ (2, 9) and TcpP overexpression is sufficient to activate P_*toxT*_ (6, 13, 14). Importantly, these same studies show that overexpression of the other TTR (ChiS or ToxR) does not activate their target promoters. These results suggest that TfoS and TcpP are sufficient to activate their target promoters, but that TTR coordination is somehow reinforced when these TTRs are expressed at native levels. However, the molecular requirements for this TTR coordination remain unclear. Here, we use ChiS/TfoS as a model system to dissect the molecular features that are required for TTR coordination.

## RESULTS

### ChiS has higher affinity for P_tfoR_ than TfoS

We have previously shown that ChiS is required to recruit the P_*tfoR*_ locus to the membrane for subsequent activation by TfoS (2). However, this recruitment is only necessary when TfoS is expressed at native levels; when TfoS is overexpressed, it bypasses the need for ChiS-dependent recruitment for P_*tfoR*_ activation (2). It remains unclear why TfoS requires ChiS-dependent membrane recruitment of P_*tfoR*_ at native levels, and why overexpression of TfoS bypasses this requirement. One hypothesis is that the TfoS DNA-binding domain (DBD) has an inherently low affinity for P_*tfoR*_, while the ChiS DBD has a higher affinity for P_*tfoR*_. This might allow ChiS to independently recruit P_*tfoR*_ to the membrane to limit its diffusion in two-dimensional space so that the promoter can be bound by the weaker DBD of TfoS.

To test this possibility, we purified the ChiS and TfoS DBDs and tested their ability to bind to P_*tfoR*_ *in vitro* through electrophoretic mobility shift assays (EMSAs). Specifically, a fluorescently labeled P_*tfoR*_ DNA probe was incubated with increasing concentrations of purified protein, and the reactions were electrophoretically separated on a polyacrylamide gel. The binding of the protein to the P_*tfoR*_ probe can be visualized by an upward shift of the fluorescently labeled P_*tfoR*_ probe on the gel (**Fig. 1A**). The percentage of the probe shifted for each sample was plotted and the line of best fit was determined by four parameter logistic (4PL) regression analysis (**Fig. 1B**). The apparent K_d_ (*i*.*e*., the amount of each protein required to shift 50% of the P_*tfoR*_ probe) was 163.5 nM for ChiS (95% CI: 144.6 nM to 183.2 nM) and 377.5 nM for TfoS (95% CI: 277.9 nM to 718.7 nM) (**Fig. 1A-B, 1D**). The steeper binding curve observed for ChiS also suggested that this protein exhibits cooperative binding to P_*tfoR*_, which is supported by a hill coefficient of >1 for the fitted curve (3.7 for ChiS vs 1.7 for TfoS). Importantly, the DNA-binding activity of the ChiS and TfoS DBDs was specific to the P_*tfoR*_ probe because they did not shift a nonspecific promoter probe (P_*VCA0053*_) at concentrations up to 750 nM (**Fig. S1**). Thus, these data indicate that ChiS may have a higher affinity for P_*tfoR*_ when compared to TfoS. An important caveat, however, is that the specific activities of these purified proteins may differ. Furthermore, the DNA binding affinities of the purified cytoplasmic domains may not match the intact TTRs *in vivo*. Western blot analysis indicates that steady state levels of ChiS are ∼5-fold higher than TfoS *in vivo* (**Fig. S2**). Also, prior work demonstrates that ChiS forms foci in the membrane, while TfoS does not (2, 8). The cooperative binding and slightly increased affinity for the ChiS DBD observed *in vitro* combined with its increased expression and propensity to form foci *in vivo* supports the hypothesis that ChiS binds to P_*tfoR*_ first, which allows for subsequent binding by TfoS to ultimately activate the promoter.

**Fig. 1.**
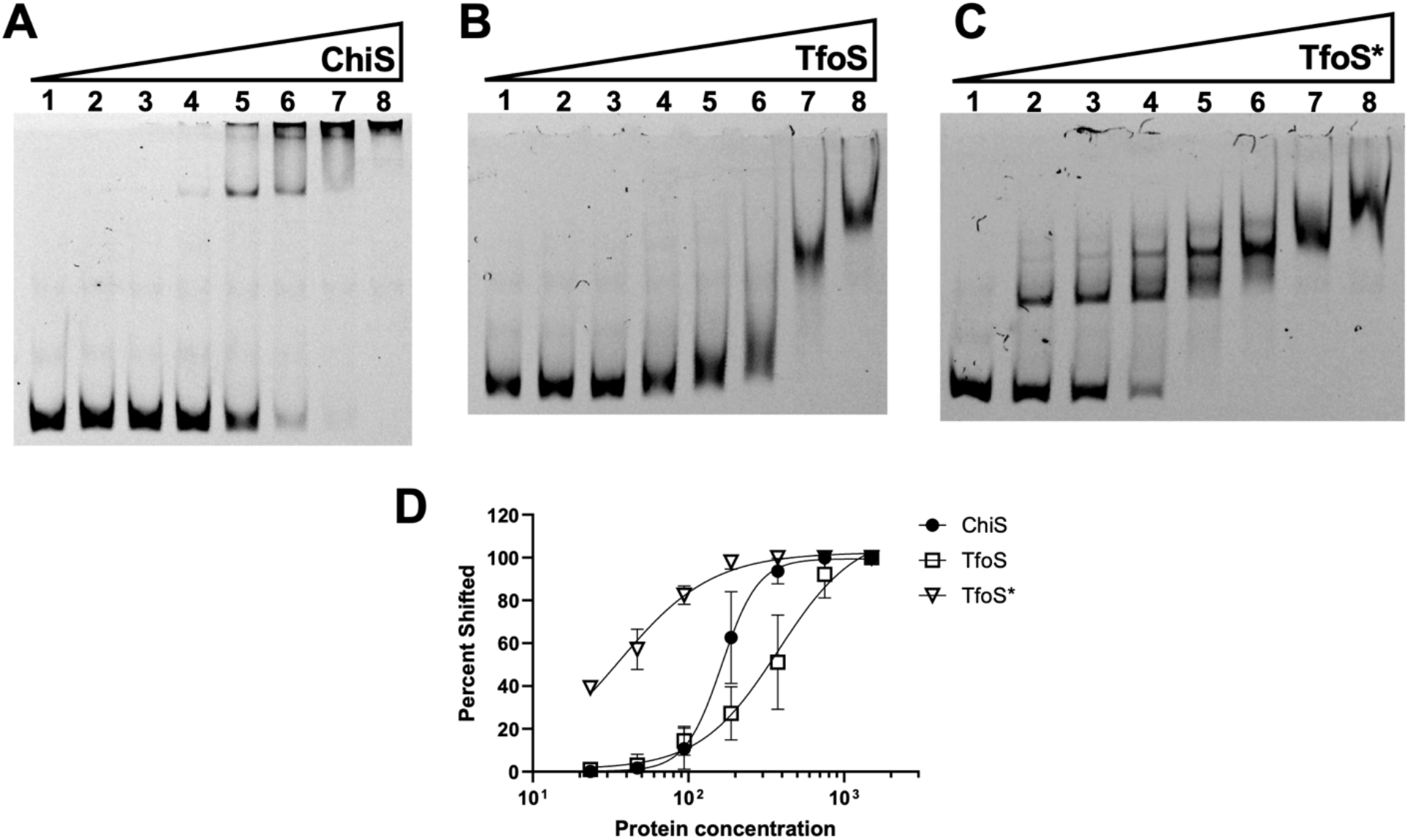
*ChiS displays a higher affinity for P*_*tfoR*_ *than TfoS*^*WT*^ *but not TfoS\**. **(A-C)** EMSAs to assess the ability of the indicated purified DBD (ChiS, TfoS, or TfoS*) to bind to the TfoR promoter (P_*tfoR*_). EMSA reactions contained increasing concentrations of the indicated protein (lane 1 = 0 nM, lane 2 = 23.4 nM, lane 3 = 46.9 nM, lane 4 = 93.8 nM, lane 5 = 187.5 nM, lane 6 = 375 nM, lane 7 = 750 nM, lane 8 = 1500 nM) and a fixed concentration (1 nM) of a Cy5-labeled P_*tfoR*_ DNA probe. Data in **A**-**C** are representative of three independent experiments. **(D)** Quantification of EMSAs shown in **A**-**C**. Lines of best fit were determined by four parameter logistic regression analysis. Data are from of three independent experiments and each data point shows the mean ± SD.

### TfoS* is a high affinity DNA-binding mutant of TfoS

If TfoS relies on ChiS-dependent membrane recruitment of P_*tfoR*_ because of its comparably weaker DBD, then we hypothesized that a TfoS mutant with a higher affinity DBD would bypass the need for ChiS. In a previous unrelated genetic screen for TfoS variants with heightened activity, we isolated an allele that contained a missense mutation within the DBD, TfoS^E1058G^ (henceforth called TfoS* for simplicity) (**Fig. S3**). We hypothesized that TfoS* might have a higher affinity DBD compared to the wildtype TfoS allele. To test this, we first purified the DBD of TfoS* (**Fig. S4**) and tested its ability to bind to P_*tfoR*_ by EMSA. We observed that TfoS* shifted the P_*tfoR*_ probe at much lower concentrations with an apparent K_d_ of 35.77 nM (95% CI: 30.96 nM to 41.25 nM), indicating that it had a higher affinity for the P_*tfoR*_ locus (**Fig. 1C-D**).Importantly, this heightened affinity is specific to the P_*tfoR*_ locus because TfoS* did not shift the nonspecific promoter probe even at concentrations up to 750 nM (**Fig. S1**).

### TfoS* allows for partial P_tfoR_ activation in the absence of ChiS-mediated recruitment

Because TfoS* has a higher affinity for P_*tfoR*_ compared to TfoS^WT^, we hypothesized that TfoS* may bypass the need for ChiS-dependent recruitment of P_*tfoR*_ for activation. To test this, we generated strains harboring a P_*tfoR*_*-gfp* reporter (2) to measure P_*tfoR*_ activation. These strains also contained a constitutively expressed *mTFP1* reporter (2) to allow for normalization of overall gene expression. Cells were incubated shaking on chitin to activate ChiS and TfoS and then imaged by epifluorescence microscopy to assess reporter activation. In order to prevent ChiS-mediated recruitment of P_*tfoR*_, the promoter was truncated to delete the ChiS-binding site (P_*tfoR*_^*ΔCBS*^) as previously described (2). This truncation prevents ChiS binding at P_*tfoR*_, which results in loss of P_*tfoR*_ activation (and therefore loss of chitin-induced natural transformation) when TfoS is expressed at native levels (**Fig. 2A-C**). Thus, P_*tfoR*_^*ΔCBS*^ allows us to assess the ability of TfoS* to activate P_*tfoR*_ in the absence of ChiS-dependent recruitment.

**Fig. 2.**
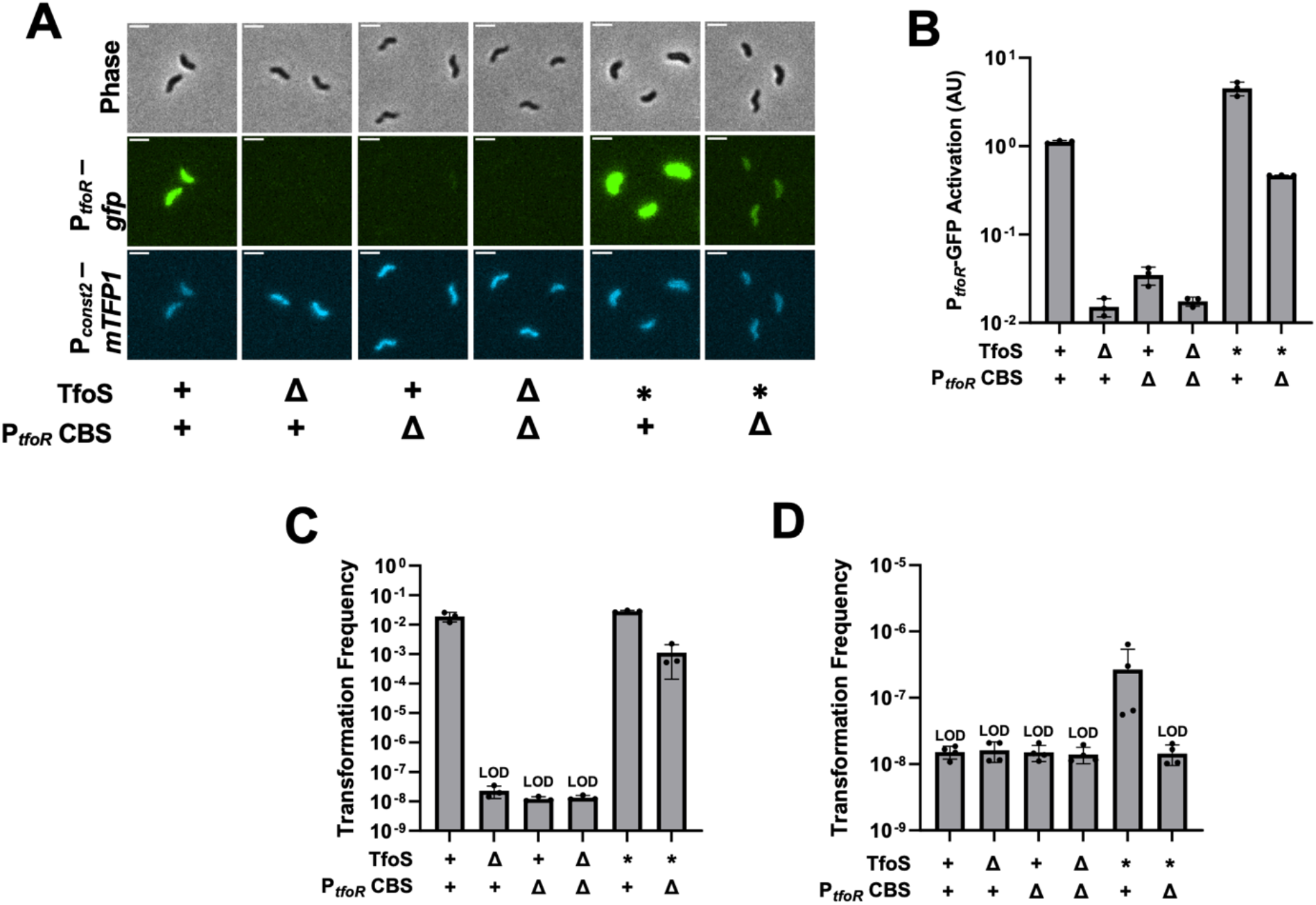
*TfoS* promotes P*_*tfoR*_ *activation in the absence of ChiS-mediated recruitment*. Strains harbored a P_*tfoR*_*-gfp* reporter, a P_*const2*_*-mTFP1* reporter, and the indicated mutations. For TfoS, “+” denotes that *tfoS* is wildtype, “Δ” denotes that *tfoS* is deleted, and “*” denotes the *tfoS*^*E1058G*^. For P_*tfoR*_ CBS, “+” denotes the wildtype promoter and “Δ” denotes the P_*tfoR*_^*ΔCBS*^promoter where the ChiS binding-site has been deleted. **(A)** Transcriptional reporter assays to assess chitin-dependent expression of P_tfoR_. Representative images of cells. Scale bar, 2µm. **(B)** Quantification of the transcriptional reporter assays depicted in **A**. For each biological replicate (*n* = 3), the geometric mean fluorescence was determined by analyzing 300 individual cells. **(C)** Chitin-dependent transformation assay of the indicated strains. Results are from three independent biological replicates and shown as the mean ± SD. **(D)** Chitin-independent transformation assays of the indicated strains. Results are from four independent biological replicates and shown as the mean ± SD. LOD, limit of detection.

We found that in the P_*tfoR*_^*ΔCBS*^background, TfoS* restored P_*tfoR*_ activation and natural transformation to almost wildtype levels (**Fig. 2A-C**). Our prior work demonstrates that overexpression of TfoS is sufficient to bypass ChiS-dependent recruitment in P_*tfoR*_^*ΔCBS*^ (2). To test whether our mutation was simply increasing TfoS levels in the cell, we compared TfoS^WT^ and TfoS* abundance by western blot. We observed that the expression of TfoS* is indistinguishable from TfoS^WT^ (**Fig. S5**), indicating that the restoration observed in TfoS* is not due to an increase in protein levels. Together, these results indicate that TfoS* bypasses the need for ChiS-dependent recruitment because it can now bind P_*tfoR*_ independently via its high-affinity DBD.

### TfoS-ChiS coordination is critical for proper regulation of natural transformation and optimal chitin catabolism

In *V. cholerae*, chitin is a required cue for ChiS-TfoS-dependent induction of natural transformation. Above, we show that TfoS* bypasses the requirement for ChiS to activate P_*tfoR*_. This result indicates that TfoS* effectively disrupts ChiS-TfoS coordination. We hypothesized that one potential consequence for disrupting ChiS-TfoS coordination would be the inappropriate induction of natural transformation in the absence of chitin. Consistent with this hypothesis, we found that TfoS* promotes a low level of natural transformation in the absence of chitin (**Fig. 2D**), suggesting that disruption of ChiS-TfoS coordination can lead to inappropriate P_*tfoR*_ activation.

The coordinated activity of ChiS and TfoS for P_*tfoR*_ activation is also critical for *V. cholerae* to grow on chitin (2, 9, 10). Thus, we hypothesized that disrupting ChiS-TfoS coordination via TfoS* would reduce *V. cholerae* fitness on chitin. To test this, we performed competitive growth assays of the parent (TfoS^WT^) vs TfoS* in both 1) the rich medium LB and 2) a minimal medium where chitin was the sole carbon source. We found that TfoS and TfoS* competed equally in LB (**Fig. 3**). However, on chitin, we found that TfoS* was outcompeted by TfoS^WT^ (**Fig. 3**). The reduced fitness observed when ChiS-TfoS coordination is disrupted suggests that this coordination is important for optimal growth on chitin.

**Fig. 3.**
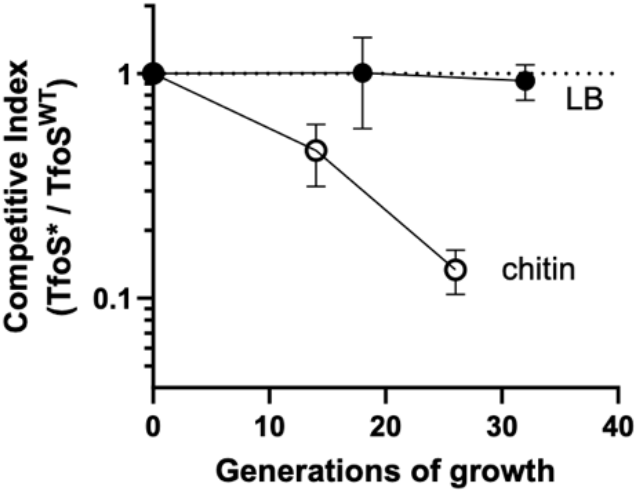
*TfoS outcompetes TfoS* during growth on chitin*. Competitive growth assays were performed between the parent (TfoS^WT^) and TfoS* in both LB medium (filled circles) and in M9 minimal media with chitin as the sole carbon source (open circles). A competitive index value of 1 indicates that strains compete equally, a value >1 indicates that TfoS* outcompetes TfoS^WT^, and a value of <1 indicates that TfoS^WT^ outcompetes TfoS*. Data are from three independent biological replicates and shown as the mean ± SD.

## DISCUSSION

Our results suggest that the cooperative activation of P_*tfoR*_ by TfoS and ChiS in wildtype cells is established by the naturally lower affinity of TfoS for DNA. Through the combination of 1) a DBD that exhibits cooperative binding and a slightly higher affinity *in vitro* and 2) a higher expression level and propensity to form foci *in vivo*, ChiS likely recruits P_*tfoR*_ to the membrane, thereby allowing the relatively lower-affinity DBD of TfoS to bind to the membrane-proximal P_*tfoR*_ locus and activate gene expression. ChiS and TfoS are likely both activated by the same inducer: chitin oligosaccharides. It is tempting to speculate that this cooperative regulatory architecture prevents the premature activation of P_*tfoR*_, ensuring that P_*tfoR*_-dependent activation only occurs when <u>both</u> ChiS and TfoS are induced by chitin oligosaccharides (*e*.*g*., as a coincidence detector (15)). Such a system might prevent chitin-associated cells from activating the potentially energetically costly behavior of natural transformation until sufficient chitin oligosaccharide nutrients have been liberated. Consistent with this model, we show that TfoS*, an allele that disrupts ChiS-TfoS coordination, results in the inappropriate induction of natural transformation in the absence of chitin. Furthermore, we show that disruption of ChiS-TfoS coordination via TfoS* impairs *V. cholerae* growth on chitin. Together, these data suggest that ChiS-TfoS coordination prevents inappropriate P_*tfoR*_ activation, and that disruption of this coordination decreases fitness under inducing conditions.

Cooperativity among TTRs has also been shown for the ToxR/TcpP system that controls virulence in *V. cholerae*. Several similarities between ToxR/TcpP and ChiS/TfoS suggest that cooperativity among ToxR/TcpP may also be established by the relative DNA-binding affinities of these TTRs. First, ToxR has been shown to have a higher affinity for P_*toxT*_ than TcpP (6). Also, overexpression of TcpP is sufficient to activate P_*toxT*_ in the absence of ToxR (12-14). These observations are consistent with ToxR-dependent recruitment of P_*toxT*_ to the membrane, which allows TcpP to bind and activate the membrane proximal promoter. Second, ToxR and TcpP are both bile-responsive transcriptional regulators (16-18). This is consistent with a regulatory architecture that could prevent the premature activation of P_*toxT*_ until both ToxR and TcpP are activated by bile. Thus, the regulatory mechanism described here for ChiS/TfoS may be a conserved feature in ToxR/TcpP. TTRs are understudied and remain completely uncharacterized in many microbes (1). Therefore, it is possible that cooperativity among TTRs via the mechanism described here for ChiS/TfoS is a broadly conserved feature among this protein family.

Our results suggest that one important feature that establishes cooperativity among TTRs is the relative affinity of their DBDs. However, this is likely not the only factor required for this cooperativity. Our results show that TfoS* has a higher affinity for P_*tfoR*_ than ChiS (**Fig. 1**); however, TfoS* does not fully activate P_*tfoR*_ and still benefits from ChiS binding within the P_*tfoR*_ promoter (**Fig. 2**, compare TfoS* P_*tfoR*_ to TfoS* P_*tfoR*_^*ΔCBS*^). Given that TfoS* has a high affinity DBD, why does ChiS binding within P_*tfoR*_ further stimulate activation? One exciting possibility is that binding of P_*tfoR*_ by both TfoS and ChiS generates a topological constraint in the bound DNA, which promotes P_*tfoR*_ activation. This topological constraint is only possible because these proteins bind to the promoter while remaining anchored in the membrane - a feature of TTRs that distinguishes them from canonical cytoplasmic transcription factors. One observation that supports this hypothesis is that ChiS activation of P_*chb*_ (a TfoS-independent promoter) requires two distinct ChiS binding sites that are helically phased (8), indicating that P_*chb*_ is likely bound by two independent ChiS dimers, which could generate a topological constraint even though it is only bound by one TTR. Furthermore, it has been shown that cytoplasmic variants of ChiS can no longer activate P_*chb*_, despite retaining the ability to bind this promoter *in vivo* (8). However, re-tethering the ChiS cytoplasmic domain to the membrane with an alternate transmembrane scaffold restores activation (2), perhaps due to restoration of topological remodeling. Future work will focus on testing whether TTR-induced topological constraints can influence the expression of their target promoters.

## METHODS

### Bacterial strains and culture conditions

All mutant strains used in this study were derived from the *Vibrio cholerae* El Tor strain E7946 (19). Strains were grown in LB medium or on LB agar unless otherwise indicated. When necessary, LB was supplemented with chloramphenicol (1 µg/mL), kanamycin (50 µg/mL), trimethoprim (10 µg/mL), spectinomycin (200 µg/mL), carbenicillin (100 µg/mL), or erythromycin (10 µg/mL), zeocin (100 µg/mL). A list of all strains can be found in **Table S1**.

### Strain construction

Mutant constructs were made using splicing-by-overlap extension (SOE) PCR exactly as previously described (20). Constructs were introduced into strains using either chitin-induced natural transformation or genetically induced natural transformation using an IPTG-inducible *tfoX* plasmid (pMMB-*tfoX*) (21). The pMMB-*tfoX* plasmid was cured before the strains were used in any experiment. Constructs were verified by PCR and/or sequencing. A list of all primers used in this study can be found in **Table S2**.

### Protein purification

The ChiS cytoplasmic domain (377-1129) was purified as an MBP-fusion as previously described (8). For purification of the TfoS and TfoS* DNA-binding domains (997-1114), proteins were tagged with GST using a pGex derivative vector. Then, *E. coli* BL21 harboring the vector of interest was grown at 37°C shaking in LB broth supplemented with 100 µg/mL carbenicillin to an OD_600_ of ∼0.6. Next, IPTG was added to the flasks to a final concentration of 1mM, and induction of protein expression was carried out at 20°C shaking for sixteen hours. Cells were harvested by centrifugation. For lysis, cells were resuspended in GST-LysB buffer (20 mM Tris-HCl pH 8.0, 250 mM NaCl, 1 mM EDTA, 2 mM DTT) supplemented with 0.4 mM PMSF, 4 mg/mL DNase, 0.4X protease inhibitor cocktail, 0.1 mg/mL lysozyme, and 1.2X Fastbreak, then incubated rocking at room temperature for twenty minutes. Cells were then lysed by sonication, clarified by centrifugation, and applied to a GSTrap HP column using an AKTA Start instrument. The column was washed with GST-LysB buffer, and then the protein was eluted via a gradient of GST-EB (100 mM Tris-HCl pH 8.0, 250 mM NaCl, 1 mM EDTA, 20 mM L-glutathione reduced). Fractions were separated on a 15% SDS-PAGE gel, and fractions containing the protein of interest were pooled. Pooled fractions were loaded into dialysis tubing and dialyzed overnight at 4°C in Storage Buffer (100 mM Tris-HCl pH 8.0, 250 mM NaCl, 1 mM EDTA, 10% glycerol). Storage buffer was replaced the next morning, and then dialysis continued in fresh storage buffer for another four hours. The dialyzed protein sample was mixed 1:1 with glycerol and stored at - 80°C until use. To verify the purity of protein preparations, samples were separated on a 15% SDS-PAGE gel. The gel was stained with Coomassie to visualize protein samples.

### Electrophoretic mobility shift assays (EMSAs)

Binding reactions contained 10 mM Tris-HCl pH 7.5, 1 mM EDTA, 10 mM KCl, 1 mM DTT, 50 µg/mL BSA, 0.1 mg/mL salmon sperm DNA, 5% glycerol, 1 nM DNA probe, and purified protein (MBP-ChiS cytoplasmic domain (8), GST-TfoS DBD, or GST-TfoS^E1058G^ DBD) at 23.4375 nM, 46.875 nM, 93.75 nM, 187.5 nM, 375 nM, 750 nM, and 1500 nM concentrations (diluted in 10 mM Tris-HCl pH 7.5, 10 mM KCl, 1 mM DTT, and 5% glycerol). Reactions were incubated at room temperature in the dark for 20 minutes and then electrophoretically separated on 0.5X Tris Borate EDTA (TBE) polyacrylamide gels at 4°C. Gels were imaged for Cy5 fluorescence on a Typhoon instrument. DNA probes were made by Phusion amplification followed by end-labeling probes with ∼20 Cy5-dCTP molecules using Terminal deoxynucleotidyl Transferase (TdT).

For quantification, analysis was conducted using the Gels feature in Fiji. Lanes were defined by a box encompassing each lane of the gel. The box size remained consistent for each lane within the same gel. Lanes were then plotted to show the intensity across the lane. The area under the curve was calculated for the unshifted and shifted band intensities, and these values were then added together to obtain the total intensity of a given lane. The shifted intensity was divided by the total intensity to determine the percent shifted.

### Chitin-induced natural transformation assays

Cultures were grown overnight in LB broth at 30°C and then subcultured and grown to an OD_600_ of 0.5-1.0. Subcultured cells were washed and resuspended in instant ocean medium (IO; 7g/L, Aquarium Systems Inc.). Then, 1 mL chitin reactions were made by mixing 10^8^ cells suspended in 100 µL IO with 150 µL chitin slurry (8g/150mL, Alfa Aesar) and 750 µL IO. Cells were incubated on chitin at 30°C static for ∼18 hours. Then, 550 µL supernatant was removed and 500 ng of transforming DNA (tDNA) was mixed with the cells by inversion. The tDNA replaced a frame-shifted transposase gene with an erythromycin resistance marker (ΔVC1807::Erm^R^). DNA and cells were incubated for five hours at 30°C static. Following incubation, 500 µL LB broth was added to the chitin reactions, and reactions were outgrown at 37°C shaking for two hours. Reactions were then plated for quantitative culture on both plain LB agar to quantify total viable counts and LB agar with erythromycin (10 µg/mL) to quantify transformants. Transformation frequency was calculated by dividing the number of transformants by the total viable counts. For reactions where no transformants were obtained, the limit of detection was calculated (*i*.*e*., the transformation frequency if a single transformant was obtained) and plotted.

### Chitin-independent natural transformation assays

Cultures were grown overnight in LB broth at 30°C and then subcultured and grown to late-log phase in LB broth supplemented with 20 mM MgCl_2_ and 10 mM CaCl_2_. Reactions were made by mixing 7 µL late-log cells with 350 µL IO. Next, 100 ng of transforming DNA (tDNA) was mixed with the cells by inversion. The tDNA replaced a frame-shifted transposase gene with a Cm^R^ resistance marker (ΔVC1807::Cm^R^). DNA and cells were incubated for ∼18 hours. Following incubation, 500 µL LB broth was added to the reactions, and reactions were outgrown at 37°C shaking for two hours. Reactions were then plated for quantitative culture on both plain LB agar to quantify total viable counts and on LB agar with chloramphenicol to quantify transformants. Transformation frequency was calculated by dividing the number of transformants by the total viable counts. For reactions where no transformants were obtained, the limit of detection was calculated (*i*.*e*., the transformation frequency if a single transformant was obtained) and plotted.

### Competitive growth assays

Strains were grown to midlog in LB medium and then washed and resuspended in M9 minimal medium lacking a carbon source. Next, TfoS^WT^ and TfoS* cultures were mixed 1:1 and ∼10^5^ cells were inoculated into either 1) 3 mL of LB broth or 2) 1mL of M9 minimal medium with chitin slurry as the sole carbon source (i.e., M9+chitin). This inoculum was also plated for quantitative culture on X-gal plates, which can distinguish the relative abundance of TfoS^WT^ (white colonies) from TfoS* (blue colonies). LB growth reactions were incubated rolling at 30°C overnight, while M9+chitin reactions were incubated shaking at 30°C for 48 hours. After the first round of growth, reactions were 1) plated for quantitative culture on X-gal containing plates to determine the number of generations of growth and the relative abundance of TfoS^WT^ and TfoS* in each reaction and 2) each reaction was back diluted into the same medium to continue competitive growth assays in a second round of growth (*i*.*e*., to increase the number of generations for each competitive growth assay). LB growth reactions were backdiluted ∼10^-6^ into fresh LB, and M9+chitin reactions were backdiluted ∼10^-5^ into fresh M9+chitin. Following the second round of growth, reactions were plated for quantitative culture on X-gal plates as described above to determine the number of generations of growth and the relative abundance of TfoS^WT^ and TfoS* in each reaction. The competitive index was defined as the ratio of TfoS* / TfoS^WT^ in the output divided by the ratio of TfoS* / TfoS^WT^ in the inoculum. This value was plotted against the approximate number of generations of growth observed in each reaction.

### Fluorescence reporter assays

Strains were grown overnight in LB broth at 30°C. Overnight cultures were washed and resuspended in IO. Then, 1 mL chitin reactions were made by mixing 10^8^ cells suspended in 100 µL IO with 150 µL chitin slurry (8g/150mL) and 750 µL IO. Cells were incubated on chitin at 30°C shaking for twenty-four hours. Following incubation, cells were dislodged from the chitin by vortexing, resuspended in IO, and placed on a glass coverslip beneath a 0.2% gelzan pad. Cells were imaged using an inverted Nikon Ti-2 microscope with a Plan Apo 60X objective lens, the appropriate filter cubes (YFP and CFP), a Hamamatsu ORCAFlash 4.0 camera, and Nikon NIS Elements imaging software. Image analysis was completed using the MicrobeJ plugin (22) in Fiji (23). The background fluorescence was determined by imaging a non-fluorescent strain, which was then subtracted from samples. Reporter fluorescence (YFP) was divided by constitutive fluorescence (CFP) to account for internal noise in gene expression. The geometric mean was then calculated from 300 individual cells for each replicate.

### Western blotting

Strains were grown overnight in LB broth at 30°C. Overnight cultures were washed and resuspended to OD_600_=100 in IO and then mixed 1:1 with 2X SDS-PAGE sample buffer (200 mM Tris pH 6.8, 25% glycerol, 4.1% SDS, 0.02% Bromophenol Blue, 5% β-mercaptoethanol). Samples were then boiled for 10 minutes. For protein visualization, 4 µL of each sample was separated on a 10% SDS-PAGE gel. Following separation, the proteins were transferred to a polyvinylidene difluoride (PVDF) membrane by electrophoresis. Membranes were blocked for 1 hour in 5% milk and then incubated overnight with rabbit polyclonal anti-FLAG (Sigma).

Membranes were then washed and incubated with anti-rabbit horseradish peroxidase-conjugated secondary antibody for three hours rocking at room temperature. Blots were washed before being developed with Pierce ECL Western blotting substrate. Blots were imaged with a ProteinSimple FluorChem R system.

### Summary Statistics

For summary statistics of all quantitative data in the manuscript, see **Dataset S1**.

## ACKNOWLEDGEMENTS

This work was supported by Grant R35GM128674 from the NIH to A.B.D.

---

**Fig. S1.**
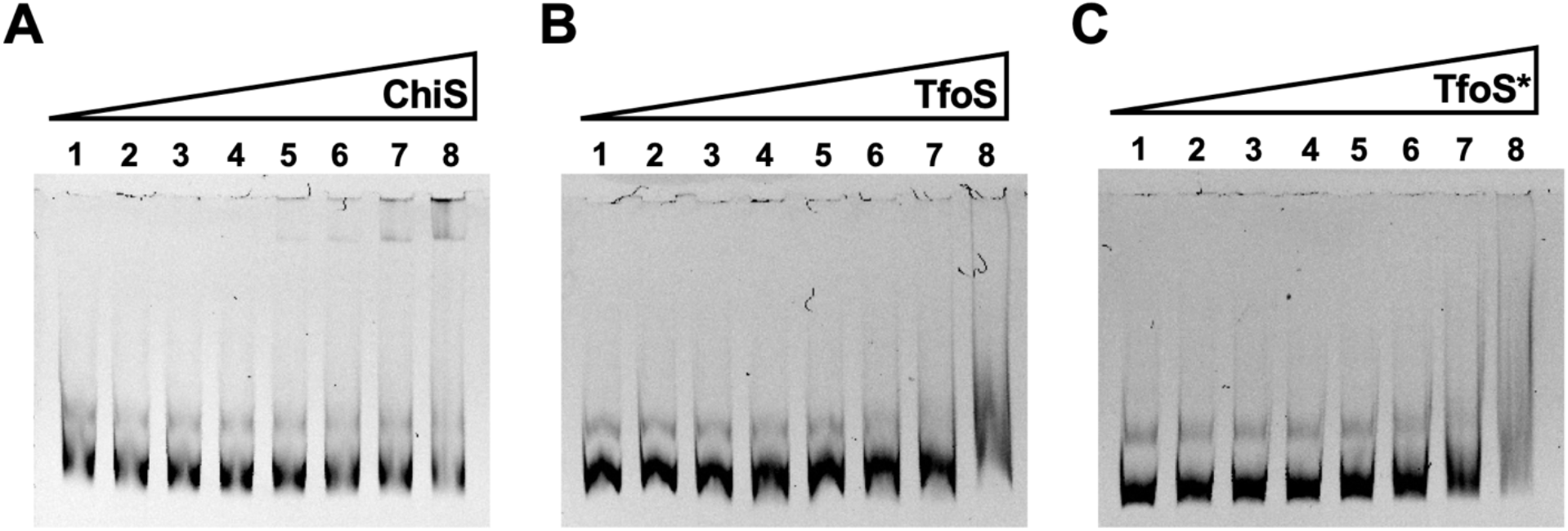
*TfoS* does not alter non-specific DNA-binding activity*. EMSAs to assess the ability of the indicated purified DBD (ChiS, TfoS, or TfoS*) to bind to P_*VCA0053*_, a promoter that is not naturally bound by any of these proteins. EMSA reactions contained increasing concentrations of the indicated protein (lane 1 = 0 nM, lane 2 = 23.4 nM, lane 3 = 46.9 nM, lane 4 = 93.8 nM, lane 5 = 187.5 nM, lane 6 = 375 nM, lane 7 = 750 nM, lane 8 = 1500 nM) and a fixed concentration (1 nM) of a Cy5-labeled P_*VCA0053*_ DNA probe. Data are representative of three independent experiments.

**Fig. S2.**
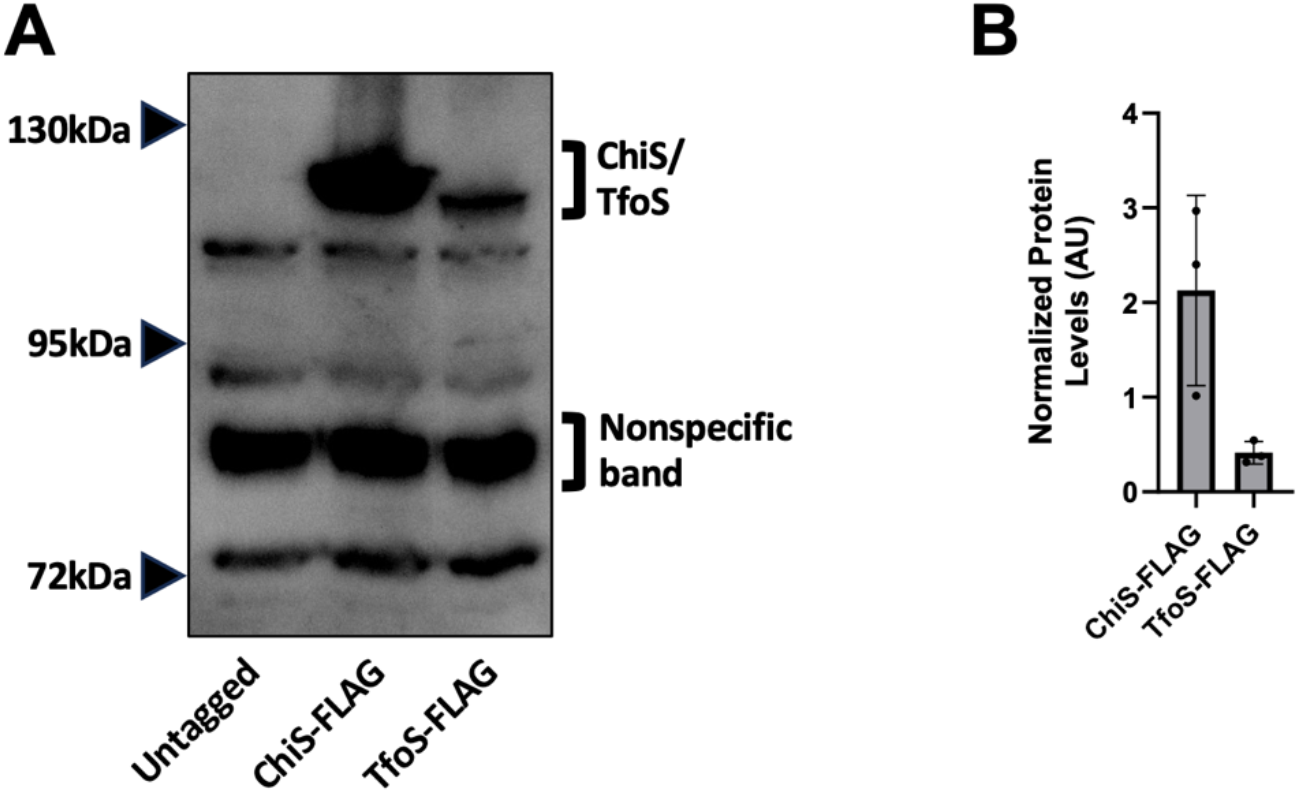
*ChiS protein levels are ∼5-fold greater than TfoS in vivo*. Western blots to assess the relative steady state levels of functional internally FLAG-tagged alleles of ChiS (ChiS^E566::FLAG^ 129 kDa; functionality demonstrated in Klancher et al. 2020) and TfoS (TfoS^N929::FLAG^ 128 kDa; see **Fig. S5** for functionality data). **(A)** A representative western blot of the indicated strains. The ChiS/TfoS band is demarcated, as well as the nonspecific band used for normalization. **(B)** Quantification of ChiS-FLAG and TfoS-FLAG levels from western blots (as in **A**). For each replicate, the normalized ChiS/TfoS level was calculated by dividing the signal intensity of the ChiS/TfoS band by the signal intensity of the nonspecific band. Data are from three independent biological replicates and shown as the mean ± SD.

**Fig. S3.**
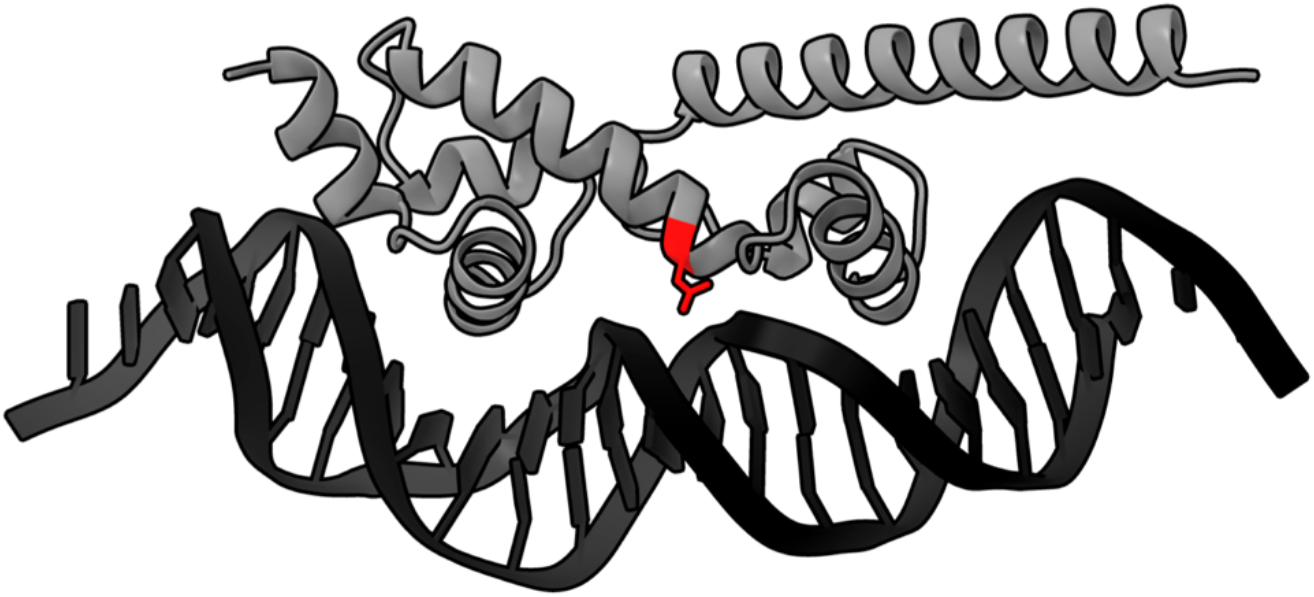
The *TfoS*^*E1058G*^ *mutation is within the predicted DNA-binding domain*. An AlphaFold3 model of the TfoS DNA-binding domain in complex with DNA. The location of the E1058 residue is shown in red.

**Fig. S4.**
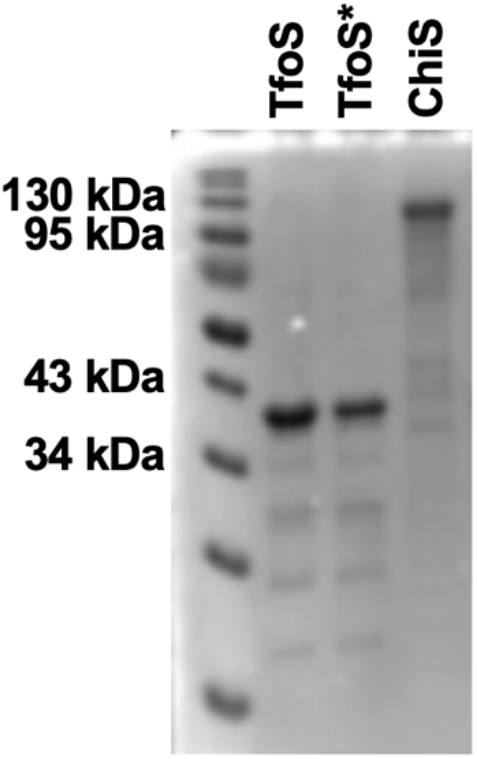
*Purified ChiS and TfoS DBDs have comparable purity*. To assess purity, 37.5 picomoles of each protein was run on a 15% polyacrylamide gel and stained with Coomassie. The expected molecular weight for TfoS/TfoS* is 40.674 kDa and for ChiS is 128.831 kDa. Data are representative of two independent experiments.

**Fig. S5.**
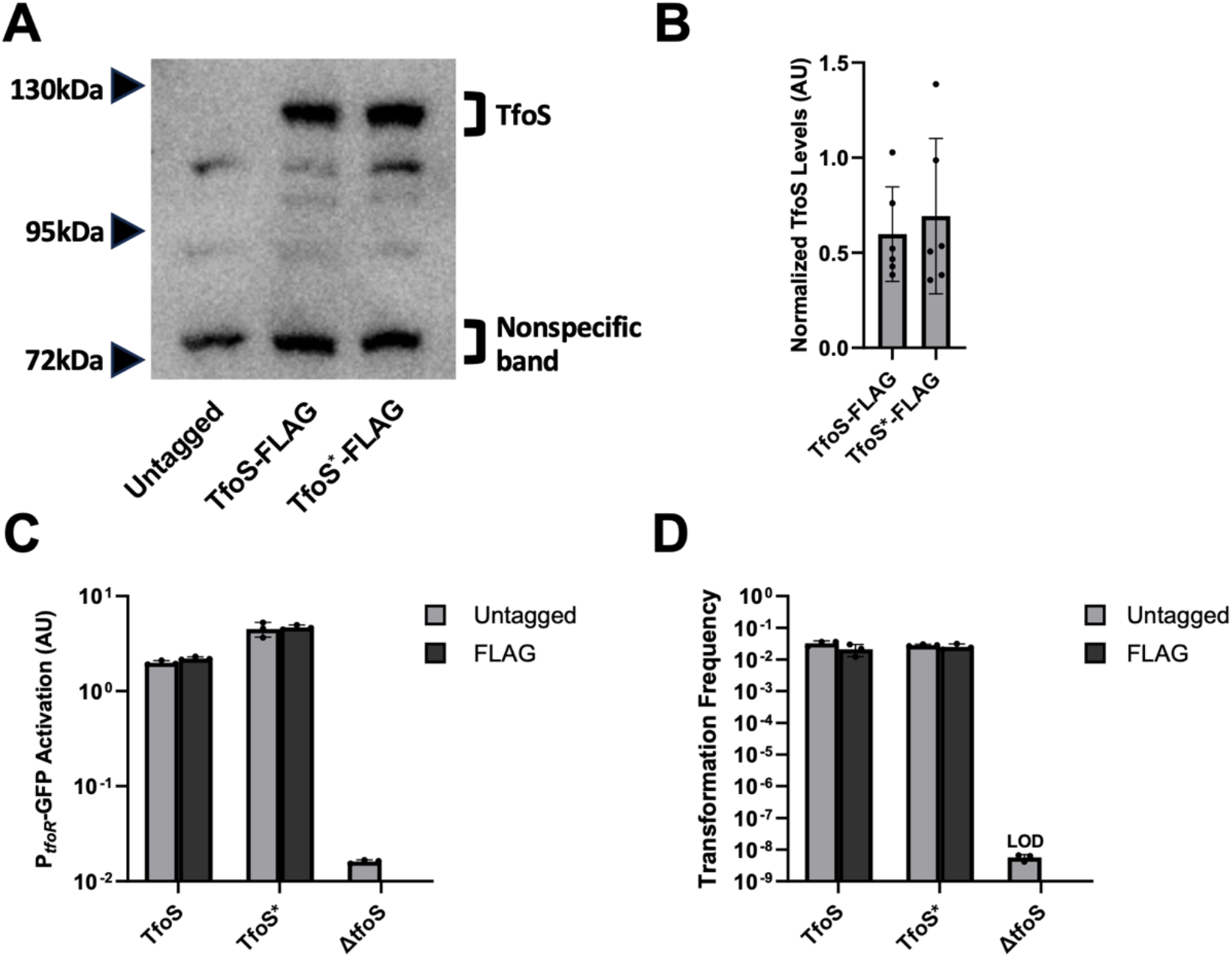
*The TfoS* mutation does not affect TfoS protein levels*. Western blots and functional activity assays were performed to assess steady state levels of TfoS and to assess the functionality of the internally FLAG tagged alleles of TfoS^N929::FLAG^ (internal FLAG tag inserted after position N929 in TfoS). **(A)** A representative western blot of strains containing the indicated mutations in the native copy of *tfoS*. The TfoS band is demarcated, as well as a nonspecific band that was used for normalization. **(B)** Quantification of TfoS levels from western blots (as in **A**). For each replicate, the normalized TfoS level was calculated by dividing the signal intensity of the TfoS band by the signal intensity of the nonspecific band. Data are from at least six biological replicates. **(C)** Transcriptional reporter assays to assess the functionality of the FLAG-tagged TfoS alleles. All strains harbored a P_*tfoR*_*-gfp* reporter, a P_*const2*_*-mTFP1* reporter, and the indicated mutations. For each replicate (*n* = 3), the geometric mean was determined by analyzing 300 individual cells. **(D)** Chitin-dependent transformation assay of the indicated strains. LOD, limit of detection. Results in **B, C**, and **D** are shown as the mean ± SD.

**Table S1.** Strains used in this study.

| Strain ID | Genotype | Reference in Manuscript |
| --- | --- | --- |
| SAD030 | *V. cholerae* WT E7946 Sm<sup>R</sup> | Parent for all *V. cholerae* strains generated in this study |
| ACH0529 / SAD4088 | *E. coli* BL21 (DE3) harboring pGex-tev TfoS DBD Carb<sup>R</sup> | Strain used for purification of TfoS DBD (residues 997-1114) |
| ACH0552 / SAD4089 | *E. coli* BL21 (DE3) harboring pGex-tev TfoS DBD<sup>E1058G</sup> Carb<sup>R</sup> | Strain used for purification of TfoS DBD* (residues 997-1114) |
| ACH0004 / SAD3634 | igVCA0265-66::Spec<sup>R</sup>-P<sub>const2</sub>-*mTFP1*; ΔVCA0692::Tm<sup>R</sup>-P<sub>tfoR</sub>-*gfp*; Δ*lacZ*::Kan<sup>R</sup>-P<sub>chb</sub>-*mCherry* | Fig. 2B-D<br>TfoS+ P<sub>tfoR</sub> CBS+ |
| ACH0014 / SAD3635 | igVCA0265-66::Spec<sup>R</sup>-P<sub>const2</sub>-*mTFP1*; ΔVCA0692::Tm<sup>R</sup>-P<sub>tfoR</sub>-*gfp*; Δ*lacZ*::Kan<sup>R</sup>-P<sub>chb</sub>-*mCherry*; Δ*tfoS*::Zeo<sup>R</sup> | Fig. 2B-D<br>Δ*tfoS* P<sub>tfoR</sub> CBS+ |
| ACH0748 / SAD4090 | igVCA0265-66::Spec<sup>R</sup>-P<sub>const2</sub>-*mTFP1*; ΔVCA0692::Tm<sup>R</sup>-P<sub>tfoR</sub><sup>ΔCBS</sup>-*gfp*; Δ*lacZ*::Kan<sup>R</sup>-P<sub>chb</sub>-*mCherry*; P<sub>tfoR</sub><sup>ΔCBS</sup> | Fig. 2B-D<br>TfoS+ ΔP<sub>tfoR</sub> CBS |
| ACH0750 / SAD4091 | igVCA0265-66::Spec<sup>R</sup>-P<sub>const2</sub>-*mTFP1*; ΔVCA0692::Tm<sup>R</sup>-P<sub>tfoR</sub><sup>ΔCBS</sup>-*gfp*; Δ*lacZ*::Kan<sup>R</sup>-P<sub>chb</sub>-*mCherry*; P<sub>tfoR</sub><sup>ΔCBS</sup>; Δ*tfoS*::Zeo<sup>R</sup> | Fig. 2B-D<br>Δ*tfoS* ΔP<sub>tfoR</sub> CBS |
| ACH0423 / SAD4092 | igVCA0265-66::Spec<sup>R</sup>-P<sub>const2</sub>-*mTFP1*; ΔVCA0692::Tm<sup>R</sup>-P<sub>tfoR</sub>-*gfp*; Δ*lacZ*::Kan<sup>R</sup>-P<sub>chb</sub>-*mCherry*; ΔVC1807::Erm<sup>R</sup>; TfoS<sup>E1058G</sup> | Fig. 2B-D ; Fig. S5C-D<br>TfoS* P<sub>tfoR</sub> CBS+ ; TfoS* Untagged |
| ACH0620 / SAD4093 | igVCA0265-66::Spec<sup>R</sup>-P<sub>const2</sub>-*mTFP1*; ΔVCA0692::Tm<sup>R</sup>-P<sub>tfoR</sub><sup>ΔCBS</sup>-*gfp*; Δ*lacZ*::Kan<sup>R</sup>-P<sub>chb</sub>-*mCherry*; P<sub>tfoR</sub><sup>ΔCBS</sup>; ΔVC1807::Erm<sup>R</sup>; TfoS<sup>E1058G</sup> | Fig. 2B-D<br>TfoS* ΔP<sub>tfoR</sub> CBS |
| TND5180 / SAD4167 | Δ*lacZ*::Spec<sup>R</sup>; igVC2075-76::Tm<sup>R</sup>-P<sub>tfoR</sub>-*gfp* | Fig. 3<br>TfoS<sup>WT</sup> |
| ACH0645 / SAD4168 | ΔVC1807::Erm<sup>R</sup>; igVC2075-76::Tm<sup>R</sup>-P<sub>tfoR</sub>-*gfp*; *tfoS*<sup>E1058G</sup> | Fig. 3<br>TfoS* |
| ACH0185 / SAD3649 | igVCA0265-66::Spec<sup>R</sup>-P<sub>const2</sub>-*mTFP1*; Δ*lacZ*::Kan<sup>R</sup>-P<sub>tfoR</sub>-*gfp*; ΔVC1807::Cm<sup>R</sup>; *tfoS* N929::FLAG | Fig. S5A-D<br>TfoS-FLAG |
| ACH0186 / SAD3650 | igVCA0265-66::Spec<sup>R</sup>-P<sub>const2</sub>-*mTFP1*; Δ*lacZ*::Kan<sup>R</sup>-P<sub>tfoR</sub>-*gfp*; ΔVC1807::Cm<sup>R</sup> | Fig. S5ACD<br>TfoS Untagged |
| ACH0510 / SAD4094 | igVCA0265-66::Spec<sup>R</sup>-P<sub>const2</sub>-*mTFP1*; ΔVCA0692::Tm<sup>R</sup>-P<sub>tfoR</sub>-*gfp*; ΔVC1807::Cm<sup>R</sup>; *tfoS*<sup>E1058G</sup> N929::FLAG | Fig. S5A-D<br>TfoS*-FLAG |
| ACH0453 / SAD4095 | igVCA0265-66::Spec<sup>R</sup>-P<sub>const2</sub>-*mTFP1*; ΔVCA0692::Tm<sup>R</sup>-P<sub>tfoR</sub>-*gfp*; Δ*tfoS*::Zeo<sup>R</sup> | Fig. S5C-D<br>Δ*tfoS* |
| CAK1295 | Δ*lacZ*::Kan<sup>R</sup>-P<sub>tfoR</sub>-*gfp* | Fig. S2<br>Untagged |
| TND2038 | Δ*lacZ*::Kan<sup>R</sup>-P<sub>tfoR</sub>-*gfp*; ΔVC1807::Cm<sup>R</sup>; *chiS* E566::FLAG | Fig. S2<br>ChiS-FLAG |
| TND1997 | Δ*lacZ*::Kan<sup>R</sup>-P<sub>tfoR</sub>-*gfp*; ΔVC1807::Cm<sup>R</sup>; *tfoS* N929::FLAG | Fig. S2<br>TfoS-FLAG |

\*Double identifiers under “Strain ID” refer to the same strain that has been stocked in two independent strain collections.

**Table S2.**
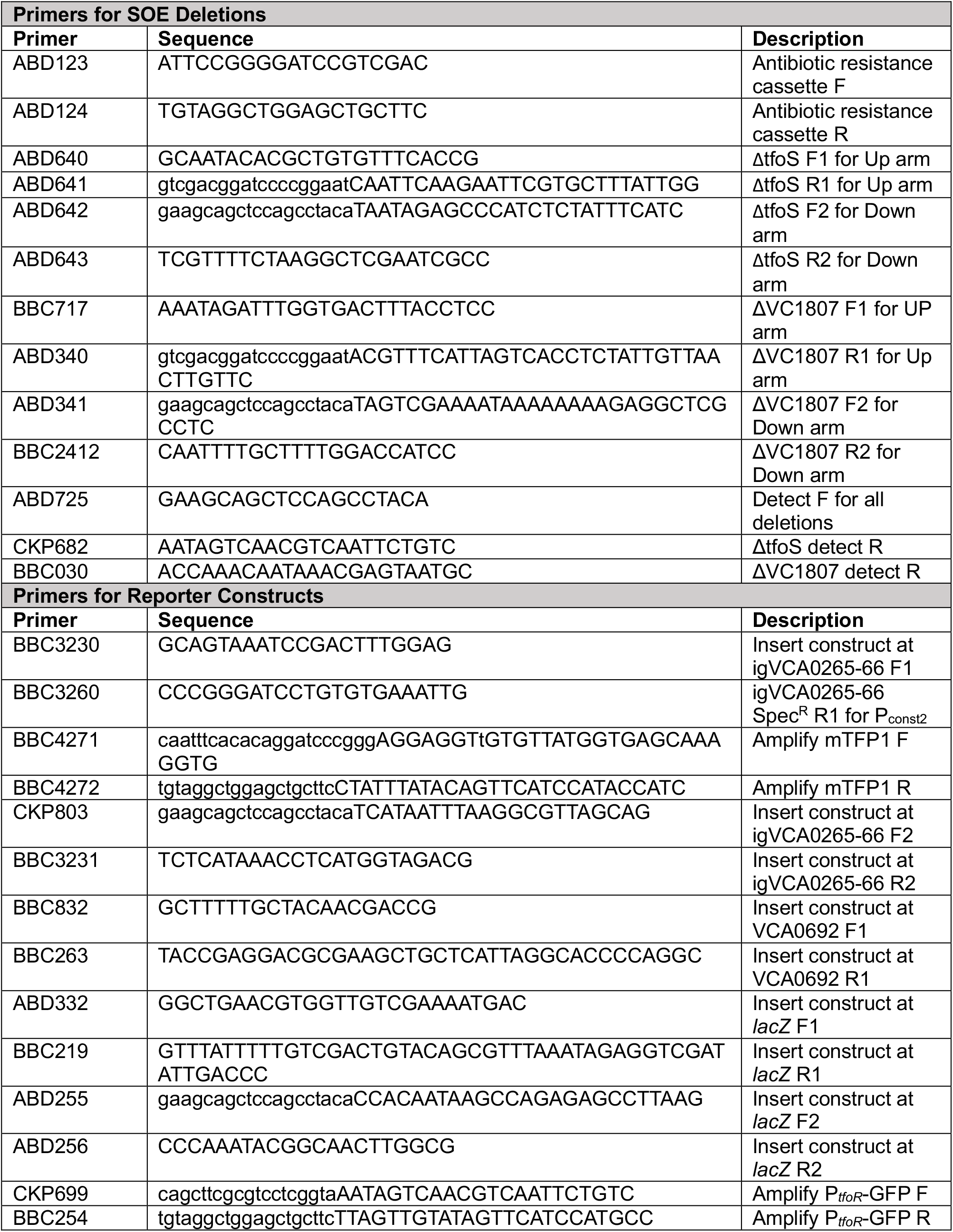
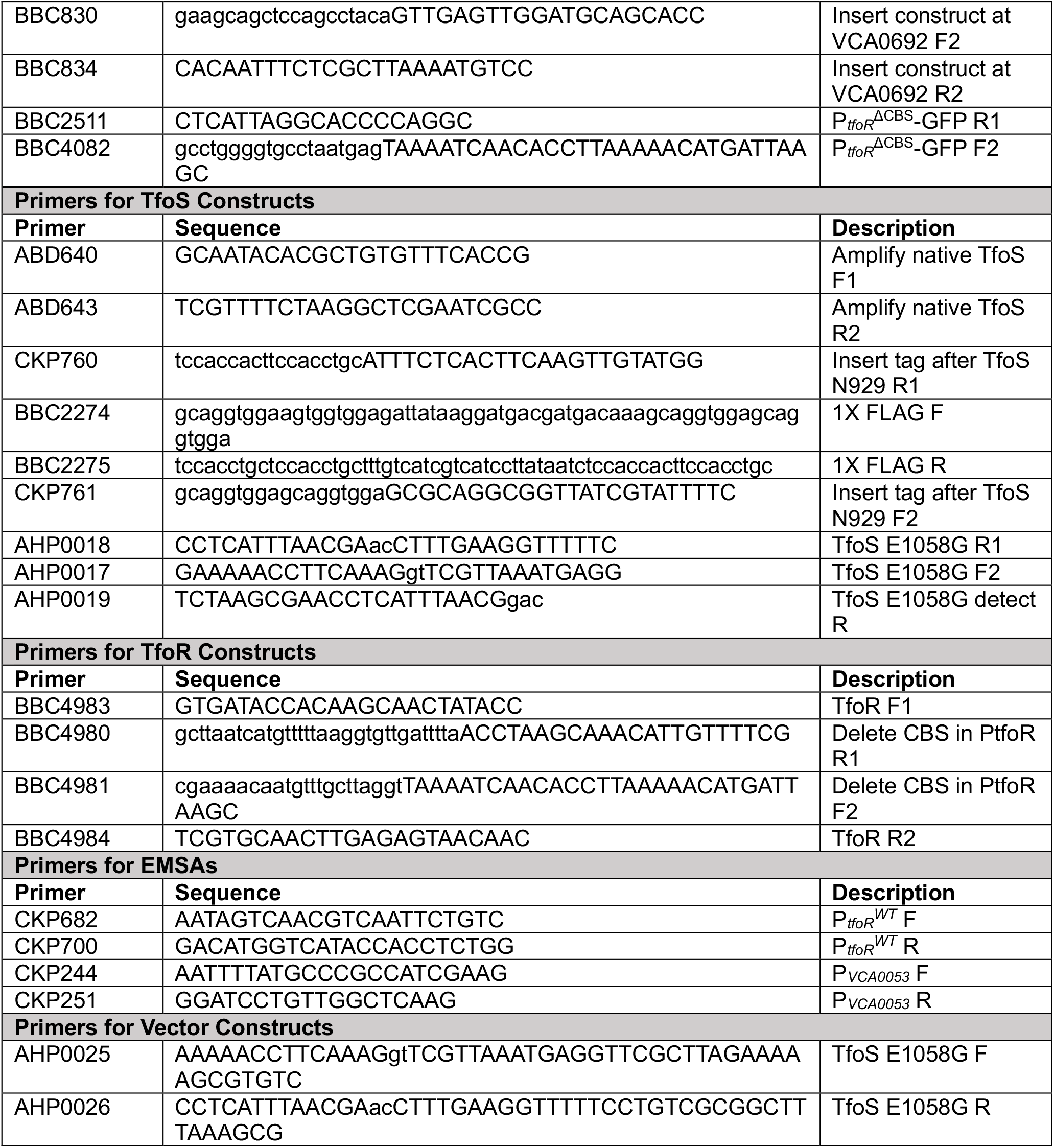
Primers used in this study.

## Notes

### Competing Interest Statement

The authors have declared no competing interest.

### Summary of Updates

A number of edits to the figures and text were made to further support the conclusions of the original submission.

